# Genetic control of RNA editing in Neurodegenerative disease

**DOI:** 10.1101/2022.08.31.505995

**Authors:** Sijia Wu, Qiuping Xue, Mengyuan Yang, Yanfei Wang, Pora Kim, Xiaobo Zhou, Liyu Huang

## Abstract

A-to-I RNA editing diversifies human transcriptome to confer its functional effects on the downstream genes or regulations, potentially involving in neurodegenerative pathogenesis. Its variabilities are attributed to multiple regulators, including the key factor of genetic variant. To comprehensively investigate the potentials of neurodegenerative disease-susceptibility variants from the view of A-to-I RNA editing, we analyzed matched genetic and transcriptomic data of 1,596 samples across nine brain tissues and whole blood from two large consortiums, Accelerating Medicines Partnership - Alzheimer’s Disease (AMP-AD) and Parkinson’s Progression Markers Initiative (PPMI). The large-scale and genome-wide identification of 95,637 RNA editing quantitative trait loci revealed the preferred genetic effects on adjacent editing events. Furthermore, to explore the underlying mechanisms of the genetic controls of A-to-I RNA editing, several top RNA binding proteins were pointed out, such as *EIF4A3, U2AF2, NOP58, FBL, NOP56*, and *DHX9*, since their regulations on multiple RNA editing events probably interfered by these genetic variants. Moreover, these variants may also contribute to the variability of other molecular phenotypes associated with RNA editing, including the functions of four proteins, expressions of 148 genes, and splicing of 417 events. All the analyses results shown in NeuroEdQTL (https://relab.xidian.edu.cn/NeuroEdQTL/) constituted a unique resource for the understanding of neurodegenerative pathogenesis from genotypes to phenotypes related to A-to-I RNA editing.

## INTRODUCTION

A-to-I RNA editing, a unique type of post-transcriptional modification, presents its diverse roles in brain development and immune regulation to contribute to the pathogenesis of neurodegenerative disease (1–3). For example, the abnormal R764G editing of the AMPA subunit, *GRIA2*, is known to affect the rate of channel desensitization and recovery from it to be involved in neurodegeneration (4). Moreover, there are other pathogenic RNA editing biomarkers proposed in previous studies, based on their aberrant editing frequencies and their functional effects on genes related to diseases (5–8). However, the current studies lack an explanation of the disease-susceptibility variants from the view of these A-to-I RNA editing biomarkers, which limits the development of drugs for treating neurodegenerative disease, although one recent study analyzed 216 Alzheimer’s blood samples for the identification of genetic variants associated with differentially edited sites (9). Due to the existence of the tissue-specific variants potentially regulating A-to-I RNA editing (10), it is necessary to provide a more comprehensive landscape for the genetic controls of RNA editing in neurodegenerative disease across different tissue types.

For this goal, we systematically studied 1,596 samples across nine brain tissues and whole blood from two large consortiums, Accelerating Medicines Partnership - Alzheimer’s Disease (AMP-AD) (11) and Parkinson’s Progression Markers Initiative (PPMI) (12,13). From the raw data of whole genomic sequencing (WGS) and RNA sequencing (RNA-seq) data, first, we identified the genetic basis of A-to-I RNA editing events among each tissue type of the neurodegenerative diseases. For a functional annotation of these variants, we then studied the potential underlying mechanisms of these associations from the view of RNA binding regulations, considering the possible effects of RNA binding proteins (RBPs) on A-to-I RNA editing (14). Moreover, we investigated the other molecular phenotypes which may be affected by these variants and also related to A-to-I RNA editing due to their possible interactions (5,6,15,16), such as protein function, gene expression, and alternative splicing, to further uncover the links and pathways between genetic variants and neurodegenerative disease. All the analysis results were archived in NeuroEdQTL (https://relab.xidian.edu.cn/NeuroEdQTL/), serving as the reference knowledgebase of the neurodegenerative pathogenesis translating genotypes to multiple phenotypes related to A-to-I RNA editing.

## MATERIALS AND METHODS

### Samples of neurodegenerative diseases

To study the genetic control of RNA editing in neurodegenerative diseases, we collected 1,050 samples of Alzheimer’s disease (AD) and 546 samples of Parkinson’s disease (PD) with both RNA-seq and WGS data from two large consortiums (Table 1), AMP-AD (11) and PPMI (12,13). The bio-specimens of these samples were obtained from whole blood, dorsolateral prefrontal cortex (DLPFC), head of caudate nucleus (AC), posterior cingulate cortex (PCC), cerebellum (CER), temporal cortex (TCX), frontal pole (FP), inferior frontal gyrus (IFG), parahippocampal gyrus (PG), and superior temporal gyrus (STG).

**Table1.** The statistics of RNA editing QTLs.

| Disease | Region | Samples | SNPs | Editing | edQTLs<sup>a</sup> |  |
| --- | --- | --- | --- | --- | --- | --- |
|  |  |  |  |  | cis | trans |
| **AD** | DLPFC | 199 | 5,863,270 | 6,791 | 38,065/24,706/917 | - |
|  | AC | 160 | 5,889,229 | 5,499 | 16,337/10,103/455 | - |
|  | PCC | 113 | 5,811,231 | 3,975 | 5,505/3,607/156 | - |
|  | CER | 78 | 5,952,311 | 35,743 | 43,687/26,444/1,395 | 2,007/3/3 |
|  | TCX | 79 | 5,969,029 | 11,384 | 7,930/5,952/262 | - |
|  | FP | 117 | 6,040,822 | 3,197 | 5,246/2,740/165 | 2,001/1/1 |
|  | IFG | 101 | 5,922,378 | 2,367 | 1,233/1,002/49 | - |
|  | PG | 97 | 6,079,696 | 2,094 | 841/651/43 | - |
|  | STG | 106 | 6,053,632 | 1,162 | 913/707/41 | - |
| **PD** | Blood | 546 | 5,767,671 | 3,617 | 91,708/42,776/1,350 | 5,578/1,805/90 |

<sup>a</sup>: The three numbers separated by forward slash were the number of pairs of RNA editing QTLs and editing events, the number of RNA editing QTLs, and the number of editing events.

### Processing of RNA-seq data

With all the samples, the first step for us was to generate individual bam files by STAR (17) using hg38 (GENCODE v.32 (18)) as the reference. These bam files were then used to detect RNA editing events by the script of REDItoolKnown.py (REDItools) (19) with default settings (e.g., minimal read coverage, 10; minimal quality score, 30; and minimal mapping quality score, 255). To ensure the confident identification of RNA editing events, we focused on known editing sites from REDIportal (January 2022) (20), removed possible SNP data from WGS and dbGap (hg38 dbsnp138 or hg19 dbsnp151) (21,22), and filtered out the candidates with supporting reads under three or editing frequency less than 0.1. Eventually, we selected one kind of RNA editing events, A-to-I RNA editing for further analysis, because of its abundance in human.

For all these detected A-to-I RNA editing events, their genome coordinates were then converted from GRCh38/hg38 to GRCh37/hg19 by LiftOver (23), to maintain the assembly consistence of RNA editing events with genotype data. Further, we selected informative A-to-I RNA editing events for quantitative trait locus (QTL) analysis, if their missing rates were smaller than 0.3 and standard deviations of editing frequencies were larger than 5% (Figure 1A). After that, there is an average of 7,583 informative A-to-I RNA editing events for each tissue group (Table 1). The editing frequencies of these events across all samples in each group were then transformed into a standard normal distribution based on rank (24), to minimize the effects of outliers on the regression scores.

**Figure 1.**
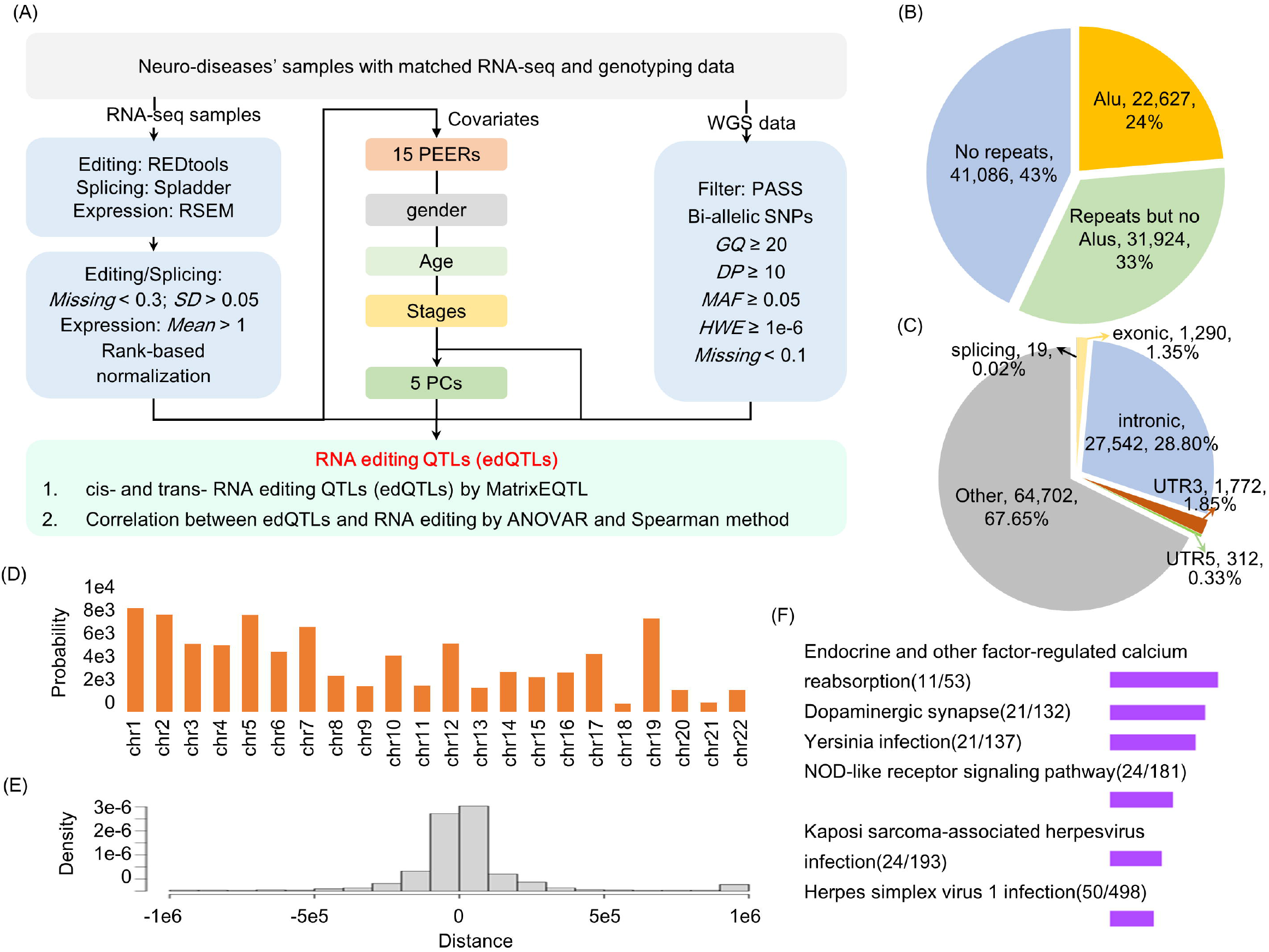
The identificaiton and distribution analysis of RNA editing QTLs. (A) The pipeline of RNA editing QTL identification. (B-D) The distributions of RNA editing QTLs in repeats, regions, and chromosomes. (E) The distances of RNA editing QTLs from the associated editing events. (F) The enriched pathways of the associated edited genes by Enrichr.

Moreover, to discover the probable interplay between genetic variants, gene expressions or splicing patterns, and A-to-I RNA editing events, we also quantified normalized read counts (TPM) by RSEM (25) and detected alternative splicing events (PSI) by SplAdder (26). Their genome coordinates and informative candidates were also processed similarly as RNA editing events above (Figure 1A).

### Processing of genotype data

The germline genotype data were directly downloaded from AMP-AD and PPMI. They were detected by standardized pipeline using BWA (27) for alignment and HaplotypeCaller mode of GATK (28) for variant calling. To increase the reliability of genotype data, bi-allelic variants were selected according to variant quality score recalibration (“PASS”), genotype quality (*GQ* ≥ 20), depth values (*DP* ≥ 10), minor allele frequency (*MAF* ≥ 5%), missing rate (*Missing* < 10%), and Hardy-Weinberg equilibrium (*P* ≥ 1 × 10^-6^). Finally, for each group, there are about 5.9 M genetic variants (Table 1) to be studied in the following RNA editing quantitative trait locus (edQTL) analysis.

### Identification of RNA editing QTLs

Like the QTL analyses performed in previous studies (29,30), we needed to consider several confounders to improve the sensitivity of edQTLs identification. For the confounders in each group, we first extracted the top five principal components (PCs) from the genotype data by smartpca (31) to correct the ethnicity differences. Next, we obtained the first 15 probabilistic estimations of RNA editing residuals (PEERs) by PEER package in R (32) to eliminate possible batch effects. These PCs and PEER factors were combined with other common confounders such as gender, age, and disease stages, as the covariates for further edQTL identification by MatrixeQTL (*FDR* < 0.05) (33).

For all these identified edQTLs, we continued to comparing the corresponding RNA editing frequencies among their three genotyping groups by ANOVA (*P* < 0.05) and also analyzing their spearman correlations with RNA editing events (*P* < 0.05). These two analyses validated the reliability of edQTL identification and also provided the visualization of the associations between genetic variants and RNA editing events.

Additionally, we also performed expression QTL (eQTL) and splicing QTL (sQTL) analysis, similarly as the pipeline of edQTLs identification. The associations between edQTLs and eQTLs or sQTLs were further explored to reveal the potentially regulatory mechanisms of these genetic variants on multiple molecular phenotypes.

### Distribution analysis of RNA editing QTLs

The functions of genetic variants are relied on their locations in specific genes or motifs. Thus, the functional annotations of these RNA editing QTLs started from the analysis of their distributions. First, we assigned edQTLs in different chromosomes, regions, and repeats, to reveal their preferred regulatory distances from corresponding editing events. Second, we identified the edQTLs potentially enriched in RNA binding proteins and their targets from chromatin immunoprecipitation sequencing (CLIPseq) analyses (34–38), since the previous studies reported their regulatory roles in A-to-I RNA editing (14,39). Third, we recognized the RNA editing QTLs in transcription factors from one previous study (40) or their altered binding motifs discovered by motifbreakR (41), for the further research about the effects of RNA editing QTLs on another phenotype changes. Last, we also analyzed RNA editing QTLs and their associated editing events in coding regions using SIFT, Polyphen2, and PROVEAN (dbNSFP version 4.2a) (42), to uncover their effects on deleterious proteins. The edQTLs in particular genes or regions may disrupt their original functions to be possibly associated with A-to-I RNA editing.

### Correlation analysis of RNA editing QTLs with disease

To uncover the potentials of RNA editing QTLs in neurodegenerative pathogenesis, we performed correlation analyses between these edQTLs and disease severity, GWAS traits, or clinical phenotypes. The severity relationships identified RNA editing QTLs possibly contributing to disease progression tested by ANOVA and Spearman method (*P* < 0.05). As for the GWAS associations, we discovered the genetic variants which were reported to be disease susceptibility loci or located in linkage disequilibrium (LD) regions with known GWAS tags using LDlinkR package (43). The parameters of this tool were set as follows: (i) *γ*^2^ (the square of the Pearson correlation coefficient of LD) threshold: 0.5, (ii) population panel: CEU (Utah residents with northern and western European ancestry), and (iii) distance limit: 500 kb. Last, the edQTLs linked with clinical phenotypes were annotated according to the archives in ClinVar (44). These correlation analyses may help pinpoint some pathogenic variants in neurodegenerative disease.

### Mediation analysis of genetic variants, editing events, and other phenotypes

To identify the shared genetic basis of A-to-I RNA editing and other phenotypes, such as gene expressions and splicing patterns, we performed mediation analysis between them using three models (45). The first model presented the genetic variants and their LD buddies (46) showing independent effects on RNA editing and other phenotypes tested by the three analyses of QTLs. The second model pointed out the potential factors acting as a bridge between genetic variants and A-to-I RNA editing (*P*_edQTL→mediators_ < 0.05, *P*_mediation effect_ < 0.05). The last model uncovered the possible downstream effects of genetic variants through the mediation of A-to-I RNA editing (*P*_eQTL/sQTL→editing_ < 0.05, *P*_mediation effect_ < 0.05). These analyses, on one hand, revealed partial mechanisms underlying the associations between genetic variants and RNA editing, on the other hand, studied further downstream effects of genetic variants associated with A-to-I RNA editing in neurodegenerative disease.

## RESULTS

### The preferred regulations of RNA editing QTLs on adjacent editing events to involve in neurodegenerative pathogenesis

Through our pipeline (Figure 1A), we detected 95,637 RNA editing QTLs associated with A-to-I RNA editing in the 1,596 samples across nine brain tissues and whole blood of neurodegenerative patients (Table 1). Almost all these edQTLs (99.92%, 95,559/95,637) were also tested to have significant correlations with corresponding RNA editing events by other two methods, ANOVA and Spearman correlation. It validated the reliability of the identified RNA editing QTLs. Of them, 98.11% (93,831/95,637) showed cis-regulatory effects on the RNA editing events, while only a small part conferred trans-regulatory effects (Table 1). The preferred regulations of RNA editing QTLs on adjacent editing events were also supported by the similar favored Alu repeats, non-coding regions, and chromosome 19 as A-to-I RNA editing and their distances from associated editing sites (Figure 1B-E).

For these RNA editing QTLs, we then performed enrichment analysis (*P* < 0.05 and *Q* < 0.05) of their associated edited genes by Enrichr (47). The enriched pathways as shown in Figure 1F revealed the probably involved neurodegeneration-related biological processes of these RNA editing QTLs. For example, dopamine is an important and prototypical neurotransmitter to affect synaptic plasticity, neuronal activity, and behavior related to neurodegeneration (48,49). Deregulated calcium signaling can lead to neurodegeneration via complex and diverse mechanisms involved in selective neuronal impairments and death (50). The proteins of the NOD-like receptors family are linked with the pathophysiology of neurodegenerative diseases (51). Thus, these RNA editing QTLs may also play important roles in these neurodegeneration-related pathways through its possible regulations on A-to-I RNA editing. Here, we selected an RNA editing QTL in the well-known neurodegenerative gene of *APOE* as an example to show its potential pathogenic roles (Figure 2).

**Figure 2.**
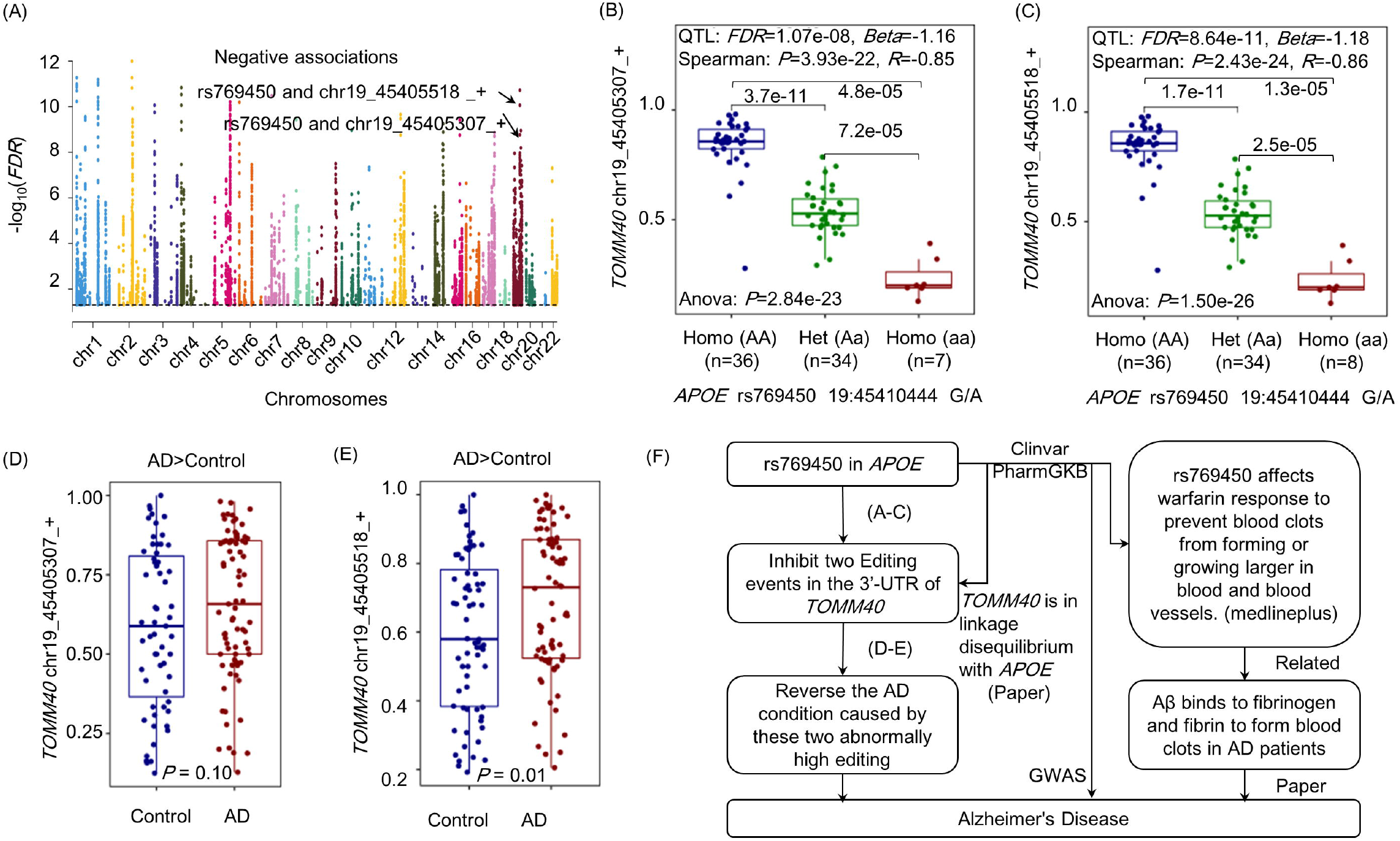
An RNA editing QTL (rs769450) showed its roles in neurodegeneration. (A-C) The Manhattan plot and boxplots showing that this edQTL negatively associated with two RNA editing events in the CER region of AD patients. (D-E) These two RNA editing events were abnormally edited in AD samples compared to controls. (F) These analyses and previous literature revealed the alleviation role of this RNA editing QTL in neurodegeneration.

This RNA editing QTL (rs769450) locates in 19:45410444 position of *APOE* gene. It showed significantly negative associations with two RNA editing events in *TOMM40* tested by all the three methods (Figure 2A-C). For these two editing events, we discovered their abnormally higher editing frequencies in AD compared to controls (Figure 2D-E), and also the potential roles of their host gene in extracellular amyloid beta aggregation (52). Thus, the down-regulation function of this RNA editing QTL on the editing events in *TOMM40* may explain its alleviation role in neurodegeneration (Figure 2F). These analyses also backed up the linkage disequilibrium relationships between *APOE* and *TOMM40* (53), the GWAS associations of this RNA editing QTL with AD traits (54), and its correlations with warfarin drug responses to prevent the formation of blood clots constituted by fibrin interacted with amyloid-β peptide (44,55).

### RNA editing QTLs may alter RNA binding regulations on A-to-I RNA editing

To explore the mechanisms of the associations between RNA editing QTLs and editing events, we analyzed that from the view of RNA binding regulations. Firstly, we identified 800 RNA editing QTLs in RBP genes (Figure 3A, Table S1). Specifically, there were 55 RNA editing QTLs in the main editing enzyme of *ADAR*, which potentially affected RNA editing events in six genes. As an example, an RNA editing QTL (rs1127311) in the 3’-UTR of *ADAR* eliminated 13 miRNA binding targets to increase *ADAR* expressions (Figure 4A-C). The altered *ADAR* expressions seemed not to regulate the editing event (chr11_65210149_+) in *NEAT1* directly (*P* = 0.997), but maybe through its binding effects on other possible editing regulators, such as *DHX9* shown in Figure 4D. The regulation of *DHX9* on the editing event was supported by their significant associations (Figure 4E) and the reported evidence of this gene to reshape global A-to-I RNA editing profiles (56). Furthermore, there were also other RNA binding proteins (Figure 4F) probably involved in this regulation. Due to the important function of *NEAT1* in the regulation of neuroglial cell mediating Aβ clearance (57,58) and the GWAS associations between this variant and the neurodegeneration-related apolipoprotein A1 levels (59), this RNA editing QTL in *ADAR* seems to be a potential biomarker in neurodegenerative disease.

**Figure 3.**
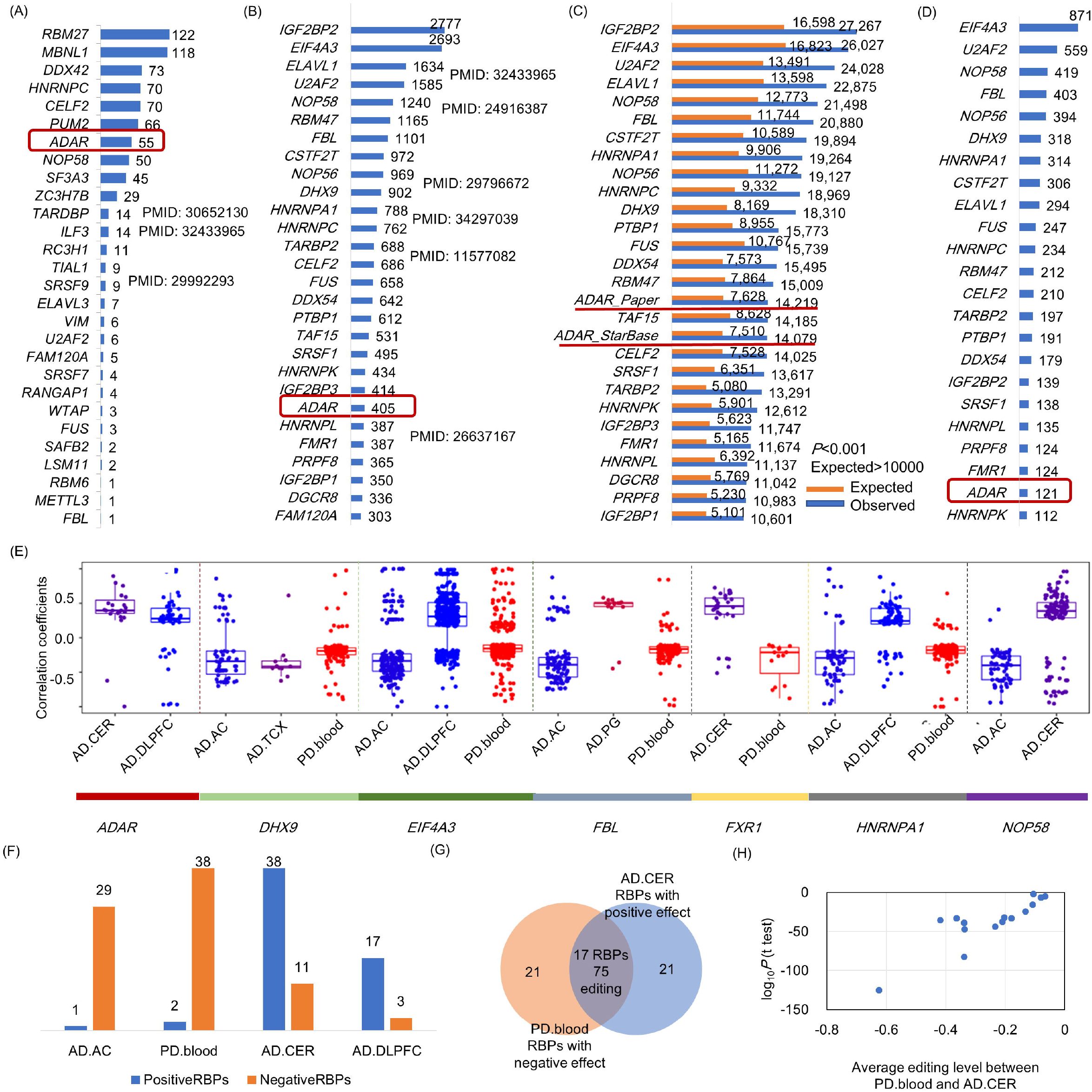
The analysis of RNA editing QTLs’ effects from the aspects of RNA binding protein (RBP) regulations. (A-B) The RNA editing QTLs (the number shown beside the bar plot) located in different RBP genes and their targets respectively. The pubmed IDs shown in the figures are the studies reporting the regulations of RNA binding proteins on A-to-I RNA editing. (C) The enrichment (expected vs. observed) of the RNA editing QTLs in RBP targets by GREGOR. The enriched ADAR binding sites from two different studies showed the similar results, revealing the reliability of the CLIPseq analyses. (D) The number of RNA editing QTLs in RBP targets potentially regulating the editing events whose frequencies were associated with the corresponding RBP expressions in the genotyping groups (AA or aa). (E) The statistically positive or negative effects of several RNA binding proteins on editing events in different groups. For others, please refer to Table S5. (F) The number of RBPs showing potentially positive or negative effects on the editing events in the four groups. For blood samples of PD and AC samples of AD, most RBPs showed negative effects on RNA editing. While in another two groups, most RBPs presented positive effects. (G-H) For the 17 RBPs showing opposite effects on 75 RNA editing events between blood samples of PD patients and CER samples of AD patients, we compared the levels of these editing events in the two groups. All the editing events were significantly down-regulated in PD, since these RBPs probably conferred negative effects in this group.

**Figure 4.**
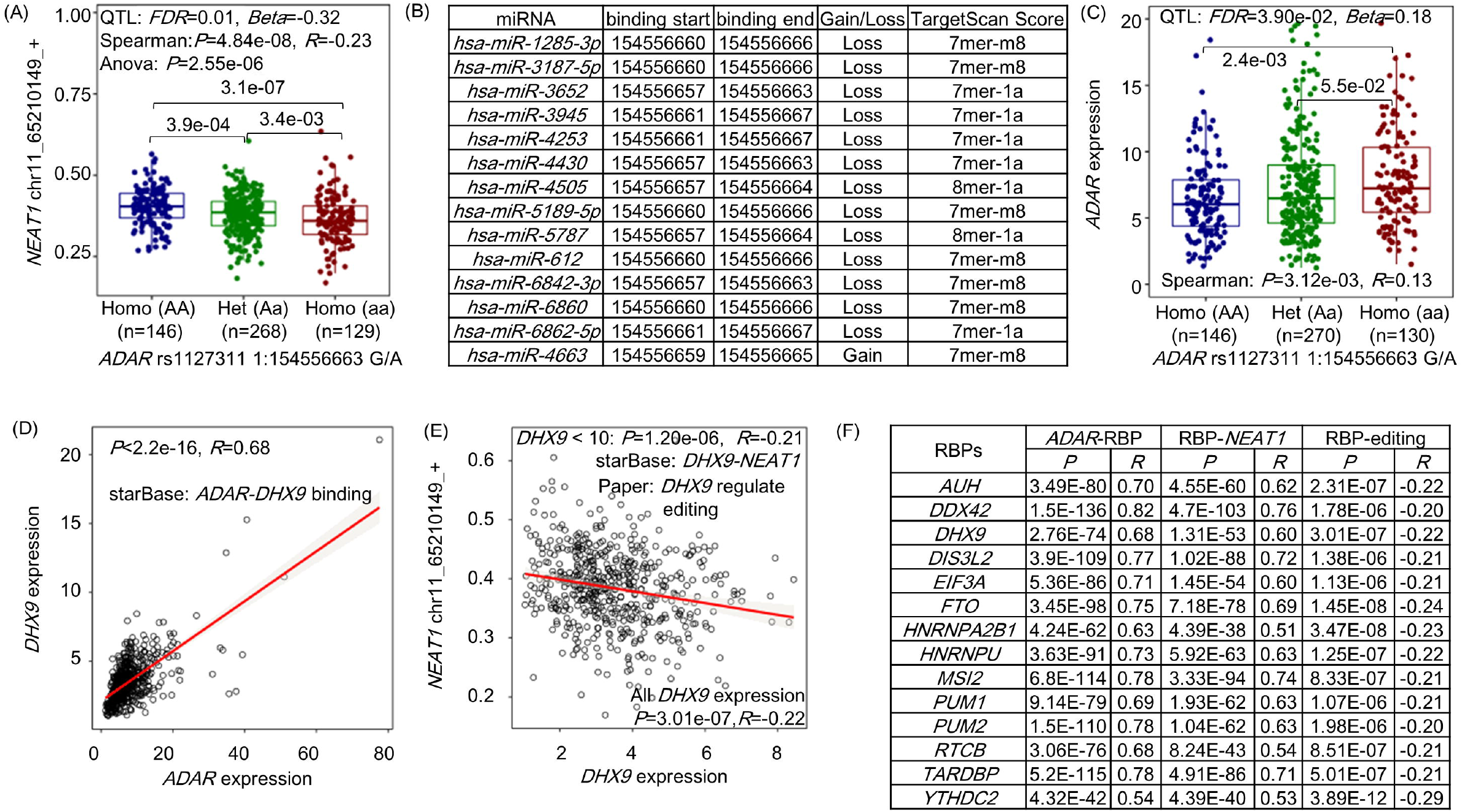
The possible effects of an RNA editing QTL on the editing event through *ADAR* regulations in the blood samples of PD patients. (A) This RNA editing QTL (rs1127311) was negatively associated with the editing event (chr11_65210149_+) in *NEAT1*. (B-C) It located in the 3’-UTR region to interfere in miRNA regulations on the gene of *ADAR*. (D) *ADAR* is an RNA binding protein to regulate other genes, such as *DHX9*. (E) The expressions of *DHX9* were also negatively associated with the frequencies of the editing event in *NEAT1*. (F) Except for *DHX9*, there were also other RNA binding proteins to be potentially involved in this regulation.

Next, we overlaid RNA editing QTLs and evaluated global enrichment of RNA editing associated variants (36) in the dataset of RBP targets from CLIPseq analyses of StarBase and one *ADAR* binding study (35), as shown in Figure 3B-C. Both analyses identified the most enriched *IGF2BP2*, followed by *EIF4A3, U2AF2, ELAVL1, DHX9*, and *ADAR*. These enriched RBPs were found to be significantly interacted with the three *ADAR* enzymes across different diseases and regions (Table S2). Some of them have been validated as *ADAR* binding partners (39,60,61). The regulations of these RBPs on the RNA editing events were evaluated by their correlations in AA and aa genotypes (Table S3-4, Figure 3D). The statistical results revealed that 93.46% (12,943/13,849) cases showed significant associations in only one genotyping group, indicating that the genetic variants may eliminate or create RBP regulations on these editing events.

Specifically, rs36118024, an RNA editing QTL in *NR2C2*, may alter the bindings and regulations of *NOP58 on the* editing event in *FGD5-AS1* through their competitions to be associated with neurodegeneration. The hypothesis was supported by the significant associations between this RNA editing QTL and editing event (Figure 5A), the possible disrupted regulations of *NOP58* on the editing event by this genetic variant (Figure 5B), the locations of this RNA editing QTL and editing event around *NOP58* binding regions detected by PAR-CLIPseq analyses (Figure 5C), and the effect of *FGD5-AS1* on PI3K/Akt signaling pathway involved in neurodegenerative pathogenesis (62,63). The above analyses revealed the possible involvements of genetic variants in RNA binding regulations to be associated with the editing events in neurodegenerative disease.

**Figure 5.**
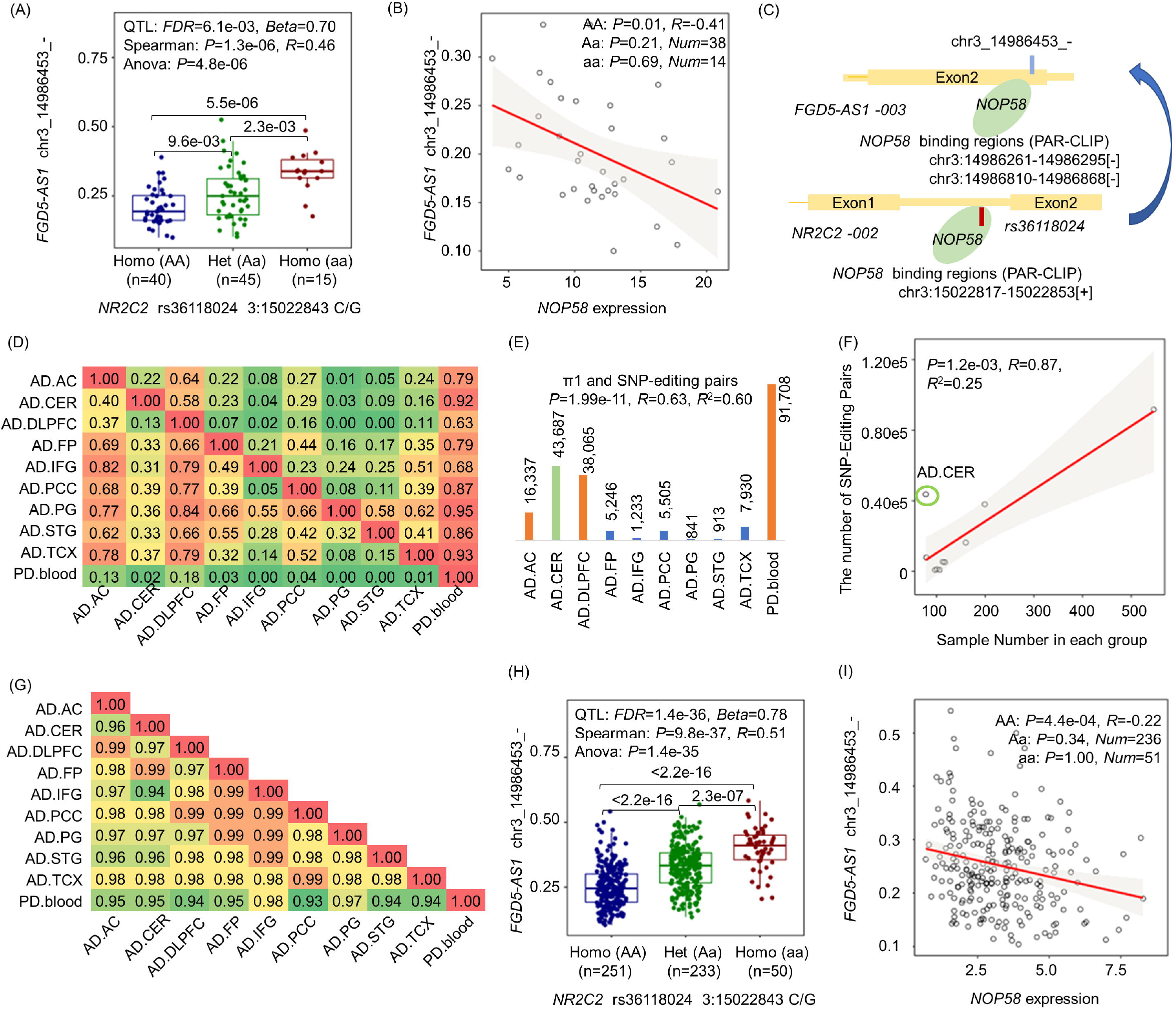
The shared genetic architecture of RNA editing across different regions and diseases. (A-C) An RNA editing QTL potentially disrupted *NOP58* regulation on an RNA editing event of *FGD5-AS1* in the STG region of AD patients. (D) π1 statistics for the significant SNP-Editing pairs in the first dataset (row) that were also shared by the second dataset (column). (E) The π1 statistics results were partially attributed to the number of significant pairs in the second dataset. (F) The SNP-Editing pairs were associated with the number of samples in each group. (G) Pearson correlation coefficients of beta values between two groups (*P* < 0.05) for the overlapped significant pairs of RNA editing QTLs and editing events. (H-I) AD and PD shared the genetic regulation of rs36118024 on the editing event in *FGD5-AS1*.

Furthermore, whether these RNA binding proteins tended to enhance or inhibit the levels of A-to-I RNA editing events was also a question needed to be addressed. To answer this, we performed one sample t-test for the significant correlation coefficients (*Num* > 10 and *P* < 0.05) between RBP expressions and the frequencies of editing events in RBP targets (Table S5, Figure 3E). Some RBPs presented consistent regulatory roles in different groups, such as *ADAR* showing positive effects as the well-known knowledge (64) and *DHX9* displaying negative effects on editing events as an RNA-independent interaction partner of ADAR (56,60). Also, additional RBPs may regulate RNA editing in bidirectional ways across different groups. For example, reported as an editing regulator (65), *FXR1* was prone to enhance editing in the cerebellum regions of AD patients and inhibit editing in the blood samples of PD patients. These differences may be attributed to the sample region heterogeneity (Figure 3F-H).

### Shared genetic architecture of RNA editing across brain regions and diseases

To test the shared genetic architecture of RNA editing across different groups, we analyzed π1 statistics (66) for the significant associations in the first dataset that were also detected in the second dataset (Figure 5D). The shared genetic architectures were partially attributed to the number of significant pairs of RNA editing QTLs and editing events in the second dataset (Figure 5E), which was associated with the number of samples (Figure 5F) similar as that of expression QTLs analysis result (*P* = 0.03, *R* = 0.69). During the analysis, cerebellum was an interesting outlier with smaller sample number, larger pairs of RNA editing QTLs and editing events, and relatively less replications from other groups. It revealed a unique genetic basis of RNA editing in cerebellum, whose functional changes in cortical and subcortical connectivity were largely disease-specific and corresponded to neurodegenerative conditions (67). Moreover, for the overlapped significant pairs, we also performed the Pearson correlation analysis of beta values between two groups. Even the pairs between AD and PD were less associated than that in different brain regions of AD, the correlation coefficients were greater than 0.93 (Figure 5G). This extremely high correlation showed the reliability of the identified RNA editing QTLs in this study and revealed the consistent effect of genetic variants on RNA editing events across different groups. Specifically, the overlapped associations of RNA editing QTLs with editing events between AD and PD may reveal the common regulatory pathways in different neurodegenerative diseases. For example, the potential regulation of rs36118024 on the editing event in *FGD5-AS1* mentioned above was also observed in the blood samples of PD patients (Figure 5H-I).

### RNA editing QTLs are potential genetic factors to modify protein functions

To uncover the effects of genetic variants on proteins, we identified 197 RNA editing QTLs which themselves or whose associated RNA editing events located in coding regions and were predicted to cause the deleterious functions of 171 proteins (Figure 6A-B). Due to the enriched functions of these proteins in the activities of microtubule and G-protein coupled serotonin receptor (Figure 6C), we could assume that the RNA editing QTLs may also involve in these neurodegeneration-related pathways (68–70). Specifically, an RNA editing QTL (rs34083116) was significantly associated with the well-known R764G editing of the AMPA subunit of *GRIA2* (Figure 6D-E). From previous studies (4,5), this editing event not only caused the deleterious protein, but also showed differential frequencies, was associated with disease progression, altered host gene expression, interfered in alternative splicing, and affected the rate of channel desensitization and recovery from it. Thus, this intergenic RNA editing QTL may also be a potential biomarker of neurodegenerative disease from its genetic effects on the well-known R/G editing event.

**Figure 6.**
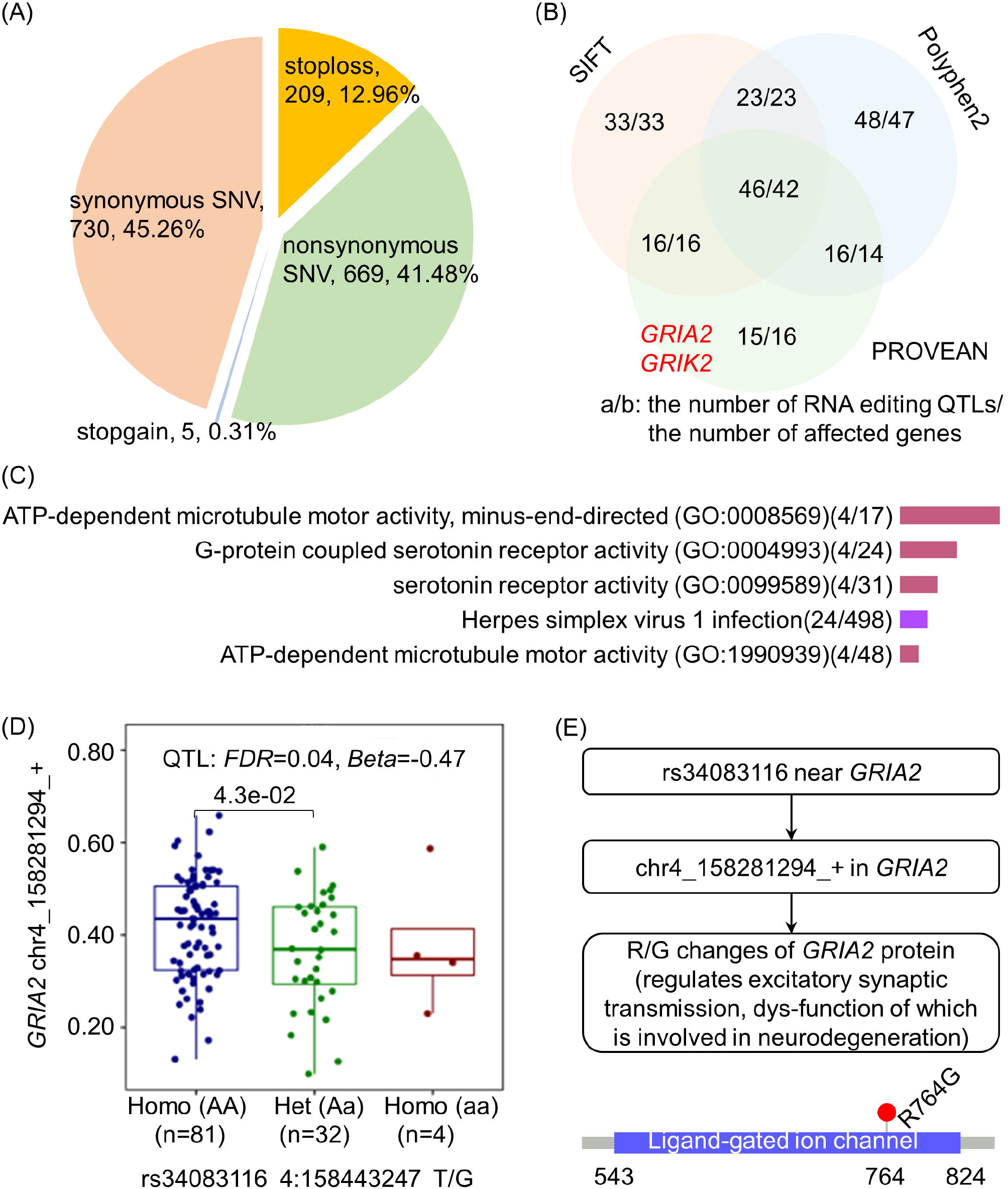
The effects of RNA editing QTLs on proteins. (A-B) RNA editing QTLs themselves or associated RNA editing events may affect amino acid sequences and protein functions. (C) The enriched biological functions and pathways of these proteins affected by RNA editing QTLs (*P* < 0.05, *Q* < 0.05). (D-E) The well-known R/G editing in *GRIA2* leading to the deleterious protein function was probably affected by rs34083116 in the FP regions of AD patients.

### RNA editing QTLs are also probable up-regulators of gene expressions

To further annotate the functions of these RNA editing QTLs, we performed mediation analysis between genetic variants, RNA editing events, and gene expressions using the models shown in Figure 7A. In total, there were 18,238 RNA editing QTLs associated with both of editing events and gene expressions (Figure 7B). Of these associations, 81.36% (47,164/57,966) showed independent genetic effects on editing events and gene expressions (model ①), 9.65% (5,594/57,966) provided propagation paths from genetic variants to A-to-I RNA editing via gene expressions (model ②), and 7.23% (4,191/57,966) explained the further effects of RNA editing QTLs on gene expressions through A-to-I RNA editing (model ③). In addition, 1.75% (1,017/57,966) associations could not be precisely classified into the second or third model of the propagations. The distributions of these associations were similar as the mediation analysis results of genetic variants, gene expressions, and epigenetic landscape of human brains (71).

**Figure 7.**
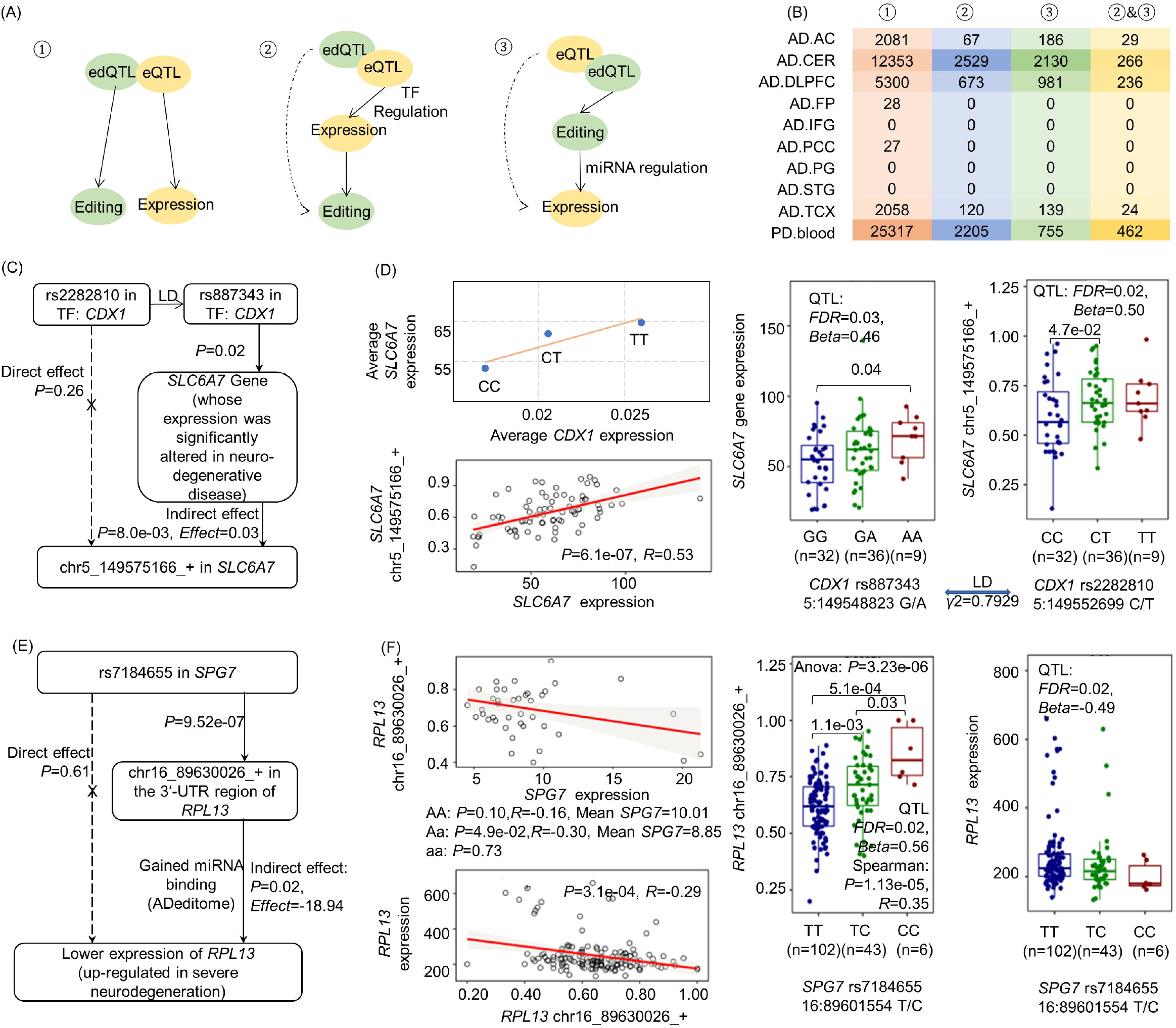
Shared genetic architecture of RNA editing and gene expressions. (A) The models showing the associations between genetic variations, RNA editing events, and gene expressions. (B) The distributions of the associations in each group. (C-D) One example in the CER region of AD patients showed that the edQTL (rs2282810) in the transcription factor of *CDX1* may affect *SLC6A7* gene expression, accompanied by the altered RNA editing event (chr5_149575166_+) in it. (E-F) One example in the AC region of AD patients showed that the eQTL (rs7184655) may affect the expression of neurodegeneration-related *RPL13*, through its regulation on an RNA editing event (chr16_89630026_+) in the 3’-UTR of this gene.

For the potential mechanisms of the second model, we explored it in the view of the transcription factor regulations. In total, there were 150 RNA editing QTLs residing in the transcription factors collected from one previous study (40). One of them (rs2282810) in *CDX1* may be associated with the regulatory functions of its LD buddy (rs887343, expression QTL, *γ*^2^ = 0.7929) on *SLC6A7* to confer its effects on the editing event. It was supported by the positive correlations of the average expressions of *CDX1* and *SLC6A7*, differential expressions of *SLC6A7* among the genotyping groups of the expression QTL, significant associations between *SLC6A7* expressions and the editing levels, and distinct editing frequencies across the genotyping groups of the RNA editing QTL (Figure 7C-D). Since *SLC6A7* belongs to SLC superfamily to play important roles in the recovery of neurotransmitters (72), the RNA editing QTL associated with the phenotype changes of this gene may be a biomarker of neurodegeneration. This example not only described the probable effects of RNA editing QTLs on gene expressions through interfering in the transcription regulations, but also revealed one possible mechanism underlying the associations between RNA editing QTLs and A-to-I RNA editing events.

To explain the natural properties of the third model, we analyzed the associations from the view of 3’-UTR regulations. In total, there were 256 RNA editing QTLs associated with the editing events in 3’-UTRs, which may interfere in miRNA regulations on the gene expressions. For example, the expression QTL analysis identified the associations between rs7184655 and *RPL13* (Figure 7E-F). This relationship may be partially attributed to the mediation of an RNA editing event in the 3’-UTR of *RPL13*. In detail, this genetic variant, also as an RNA editing QTL, probably up-regulated the 3’-UTR editing event. According to a previous ADeditome study, this editing event has been reported to create several new miRNA binding targets, thus enhancing the degenerations of *RPL13*. Due to the important roles of *RPL13* in tau interactions (73,74) and the GWAS associations between this variant and creatinine levels (75), we may annotate the functions of this RNA editing QTL in neurodegeneration. This example showed the potential mechanisms of the genetic effects on gene expressions through the mediations of A-to-I RNA editing, and also pointed out multiple phenotype changes probably caused by RNA editing QTLs in neurodegenerative disease.

### RNA editing QTLs may involve in the regulations of alternative splicing

Similarly, for the effects of RNA editing QTLs on splicing, we also performed mediation analysis between genetic variants, RNA editing events, and alternative splicing patterns. After that, we discovered 17,101 RNA editing QTLs to be associated with both of editing and alternative splicing events (Figure 8A-B). Beside the 77.94% (69,627/89,335) independent associations (model ①) and 1.58% (1,411/89,335) unclassified relationships, 6.96% (6,215/89,335) may describe the genetic effects on spicing events as the potential up-regulators of A-to-I RNA editing (model ②), and 13.52% (12,082/89,335) presented the probable associations between genetic variants and splicing events through partial mediation of A-to-I RNA editing (model ③).

**Figure 8.**
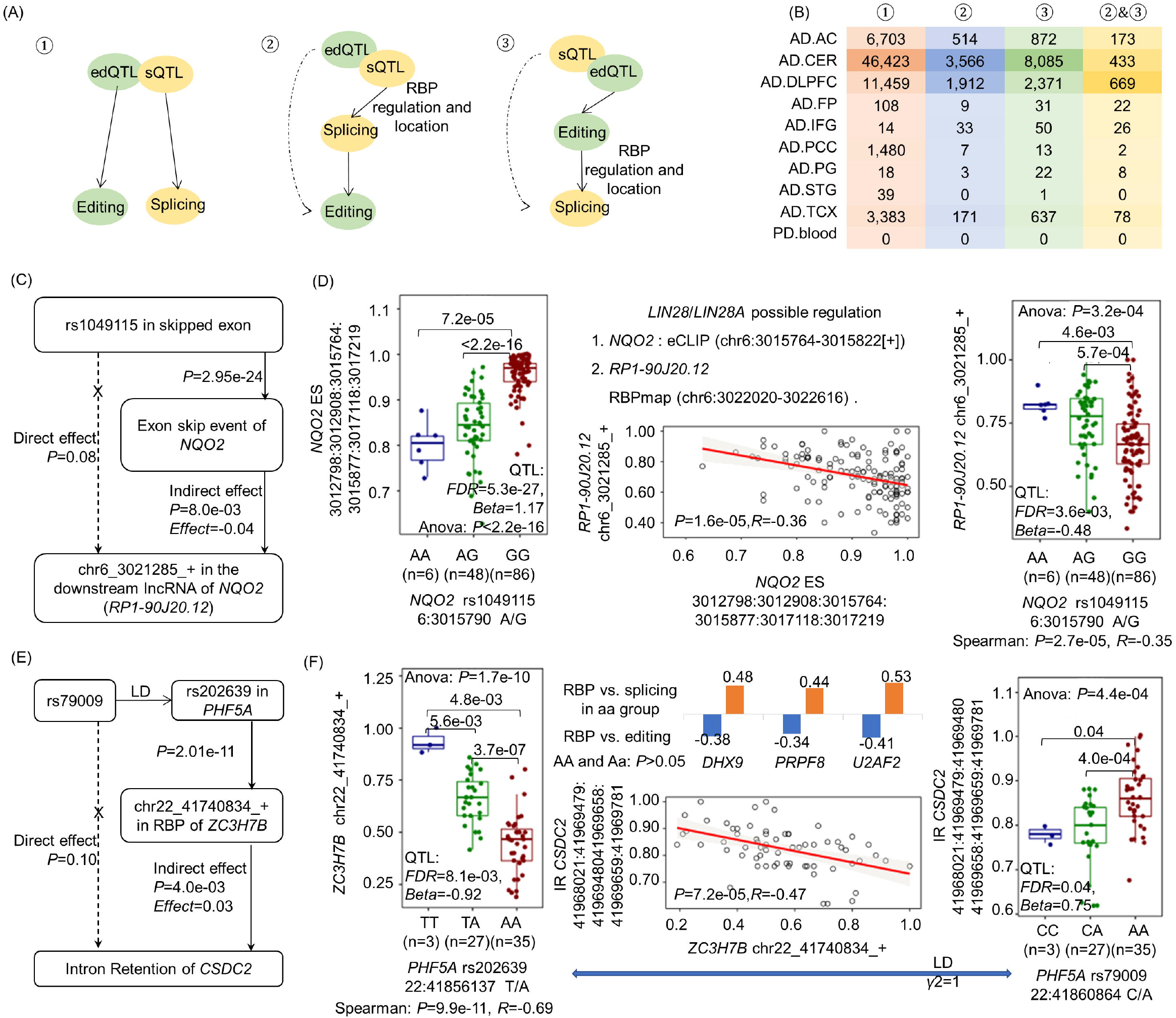
Shared genetic architecture of RNA editing and alternative splicing. (A) The models showing the associations between genetic variants, RNA editing events, and alternative splicing patterns. (B) The distributions of the associations in each group. (C-D) One example in the DLPFC region of AD patients showed that the edQTL (rs1049115) may disturb *LIN28* regulations on the exon skipping and editing event in adjacent two genes. (E-F) One example in the CER region of AD patients showed that the eQTL (rs202639) was associated with an intron retention event though partial mediation of an RNA editing event in *ZC3H7B*. For the potential mechanisms of the associations between editing and splicing, we explored the possible regulations of three RNA binding proteins on these two events, supported by their significant associations (Pearson correlation coefficients).

For the effects of RNA editing QTLs on alternative splicing in the second model, we explored the possible mechanisms from the edQTLs themselves in RNA binding proteins or splicing regions (spliced exons ± 20nt) to possibly involve in splicing regulations. After that, we identified 46 RNA editing QTLs potentially associated with alternative splicing events which were partially responsible for A-to-I RNA editing. For example, an RNA editing QTL (rs1049115), also a splicing QTL, located in the skipped exon of *NQO2* to potentially affect the regulation of *LIN28* on the splicing event (Figure 8C-D). It was supported by the differential splicing among the three genotyping groups, the eCLIP analysis for the binding of *LIN28* on the mutated region, and the regulatory roles of *LIN28* on splicing (76). Moreover, the altered splicing event seems to significantly interact with the editing event in an adjacent downstream lncRNA of *NQO2*. This lncRNA was also predicted to be a potential target of *LIN28* by RBPmap (77), and its editing event had differential frequencies among the three genotyping groups. The analyses above revealed that this genetic variant may disrupt *LIN28* regulations on the splicing and editing events of the two adjacent genes. One gene of them (*NQO2*) has showed its ability in neurodegenerative pathogenesis due to its roles in the increase of free radicals production under enhanced generation of quinone derivatives of catecholamines (78). Thus, the RNA editing QTL causing the phenotype changes of this gene could be pointed out as a potential biomarker of neurodegeneration.

For the possible mechanisms of the third model, we identified 244 RNA editing QTLs whose associated editing events located in RNA binding proteins or splicing regions to probably interfere in the splicing of exons. Specifically, a genetic variant probably affected the intron retention of *CSDC2* through partial mediation of an RNA editing event in *ZC3H7B* (Figure 8E-F). The associations between this editing and splicing event may be attributed to the co-regulations of RNA binding proteins, such as *DHX9*, *PRPF8* and *U2AF2*. Their involvements in alternative splicing and RNA editing have been showed in previous studies (39,60,79,80). According to these analyses, we could further expand the functions of RNA editing QTLs from diverse aspects of phenotype changes.

## DISCUSSION

This study aims to annotate genetic variants in neurodegenerative disease from the aspect of A-to-I RNA editing. Based on two large datasets of neurodegenerative samples, we identified 95,637 genetic variants associated with A-to-I RNA editing in total. Most of these associations belong to a kind of cis-regulatory controls, revealing the preferred regulations of genetic variants on adjacent A-to-I RNA editing (Figure 1-2). To further explore the functions of these genetic variants, we performed analyses from the following two parts. One section was to uncover the potential mechanisms for the associations between genetic variants and RNA editing events, including the interference of DNA mutations in the editing regulations by RNA binding proteins (Figure 3-5, 8) and transcription factors (Figure 7C-D). Another section was to identify other molecular phenotypes beside RNA editing that these genetic variants may affect in neurodegenerative disease. The analyses recognized genetic effects on protein recoding (Figure 6), gene expression (Figure 7E-F), and alternative splicing (Figure 8E-F) through A-to-I RNA editing. All the analyses results provided a reference knowledgebase of the neurodegenerative pathogenesis translating genotypes to multiple phenotypes related to A-to-I RNA editing.

The latter functional annotations of the genetic variants relied on the accurate identification of RNA editing QTLs. To address this, our study also performed another two analyses, ANOVA and Spearman correlation. It showed that 95.95% (216,942/217,051) associations could also pass these two tests, revealing the reliability of our QTL identification pipeline. Moreover, we also compared our pipeline with another procedure used in one previous study (71). The comparison estimated π1 statistics as 0.74 (*FDR* < 0.05) for the significant associations between genetic variants and gene expressions among DLPFC dataset in our study which were also detected using their method. It revealed the rigorousness and reliability of our pipeline. The accurate identification of phenotypes quantitative trait loci laid the foundation of further study for the functional annotations of genetic variants.

In this study, we also compared RNA editing QTLs across tissues and diseases (Figure 5D). The differences between nine brain regions of AD patients revealed the contributions of tissues to RNA editing regulation diversity (Figure 5D). This was also validated in another healthy human tissues from Genotype-Tissue Expression (GTEx) project (10). As shown in Figure S1A, two similar brain regions, cerebellar hemisphere and cerebellum, owned more association pairs of genetic variants and RNA editing events. Specifically, for cerebellum and blood samples, this hypothesis of tissue contributions also held up with more than three folds of overlapped pairs between diseases and healthy controls in the same tissue when comparing to that in different tissues (Figure S1B). Besides, the disease was also a potential factor to affect the genetic controls of RNA editing. For example, the healthy cerebellum and blood samples shared 3,255 pairs of genetic variants and RNA editing events. This number reduced for the cerebellum regions of AD patients and blood samples of healthy controls or PD patients. Interesting, compared to healthy controls, blood samples of PD patients had more overlapped pairs with cerebellum regions of AD patients (Figure S1C). It revealed the similar pathogenesis between Alzheimer’s and Parkinson’s diseases.

Due to the tissue contributions to RNA editing regulations, our work was complementary to one recent study. It identified RNA editing QTLs in peripheral blood of 216 Alzheimer’s cases with African American or White race (9), while our study recognized RNA editing QTLs in nine brain regions of AD patients and whole blood samples of PD patients. Both studies identified the roles of an RNA editing QTL (rs4723537) in neurodegeneration (Figure S2A), such as its associations with differentially edited site between AD and controls (9), and pathogenesis biomarker of tau levels (Figure S2B-C). These two studies constituted a comprehensive landscape for the genetic architecture of A-to-I RNA editing in neurodegenerative disease.

Furthermore, to uncover the potential mechanisms for the associations between genetic variants and RNA editing events, we mainly explored it from the view of interfered RNA binding effects (Figure 3), due to the possibility of allele-specific binding and editing regulations of RNA binding proteins proposed in previous studies (14,81). Beside the main editing enzyme of *ADAR*, we also discovered multiple other factors involved in the regulations of A-to-I RNA editing, including the top RNA binding proteins such as *EIF4A3, U2AF2, NOP58, FBL, NOP56*, and *DHX9*. They were all significantly associated with *ADAR* expressions (Table S2) and correlated with the editing events in their binding targets (Table S3). For some of these RBPs, previous evidence have verified the bidirectional roles of *DHX9* on RNA editing in cancers (56) and the alteration of most known editing sites in a single direction by *U2AF2* in K562 cells (39). The other RBPs can be further studied as novel RNA editing regulators, either through their interactions with main editing enzymes indirectly or binding to the adjacent editing regions directly. This is our further study to explore the potential functions and regulation pathways of the top RNA binding proteins on A-to-I RNA editing.

Last, to reveal the diverse functions of RNA editing QTLs, we also identified the other molecular phenotypes possibly affected by genetic variants and related to A-to-I RNA editing using mediation analysis (Figure 7-8). In total, 148 genes and 417 splicing events were involved in this kind of propagation paths (model ② and ③). These analyses mainly focused on the associations between single variant and each phenotype. According to the mediation results shown in NeuroEdQTL database, multiple genetic variants may confer their co-regulations on the downstream phenotypes, such as the effects of three RNA editing QTLs (rs2282810, rs2282812, and rs2240783) on the expressions of *SLC6A7* to be associated with the RNA editing event (Figure 7C-D), and the potential regulations of five RNA editing QTLs (rs7184655, rs35878298, rs77497623, rs2280370 and rs174035) on the expressions of *RPL13* through A-to-I RNA editing (Figure 7E-F). Given this possibility and the interactions between multiple editing events or genes (5), we plan to propose a multi-layer network of DNA mutation, RNA editing, and gene expression in the future, to systematically explain the potential mechanisms related to these three kinds of features in neurodegenerative pathogenesis.

## CONCLUSION

This study systematically annotates the potentials of genetic variants from the aspect of A-to-I RNA editing across nine brain tissues and whole blood of neurodegenerative disease. Specifically, it provided a reliable list of genetic variants associated with A-to-I RNA editing in two large neurodegeneration-related consortiums. Next, it tried to explain the potential mechanisms for the variant effects on editing events, revealing the top RNA binding proteins probably regulating A-to-I RNA editing. Last, it proposed three other molecular phenotypes which were affected by the genetic variants and also related to A-to-I RNA editing. The whole work, combined with one recent RNA editing QTL study in Alzheimer’s disease, will be a comprehensive and unique resource for the genetic controls of A-to-I RNA editing in neurodegenerative disease.

## ACKNOWLEDGMENTS

This work was supported by the National Natural Science Foundation of China (Grant No. 62002270), the Fundamental Research Funds for the Central Universities, the Natural Science Foundation of Shaanxi Province of China (Grant No. 2020JQ-332), the China Postdoctoral Science Foundation (Grant No. 2018M643583), National Key R&D Program of China (Grant No. 2017YFA0205202), and partially funded by the National Natural Science Foundation of China (Grant No. 61672422). The funders had no role in study design, data collection and analysis, decision to publish or preparation of the manuscript.

## SUPPLEMENTARY FIGURE LEGENDS

**Figure S1. The comparison results of RNA editing QTLs across tissues and diseases.** (A) The overlapped significant pairs of genetic variants and RNA editing events (*P* < 0.0005) between healthy samples from different tissues. (B) The overlapped significant pairs between healthy and neurodegenerative tissues. (C) The overlapped significant pairs between neurodegenerative samples from different tissues.

**Figure S2. The comparison of one recent RNA editing QTL study in the blood samples of AD patients with this study.** (A) The annotation of an RNA editing QTL in both datasets. (B) The associations of this RNA editing QTL with the editing event in this study. (C) This RNA editing QTL was significantly associated with tau levels, one biomarker of neurodegeneration.

**Table S1. Number of RNA editing QTLs in RBP genes and their targets from StarBase**

**Table S2. The correlations between RNA binding proteins and three ADAR enzymes across different tissues of neurodegenerative patients**

**TableS3. The correlations between RBPs and editing events probably affected by RNA editing QTLs in RBP targets (Mean expression > 1)**

**TableS4. The number of editing events (affected by RNA editing QTLs in RBP targets) associated with RBP expressions**

**TableS5. The potentially regulatory direction of RBPs on A-to-I RNA editing (significant associations more than 10)**

## REFERENCE

1. Behm, M. and Öhman, M. (2016) RNA editing: a contributor to neuronal dynamics in the mammalian brain. Trends Genet., 32, 165–175.

2. Gardner, O.K., Wang, L., Van Booven, D., Whitehead, P.L., Hamilton-Nelson, K.L., Adams, L.D., Starks, T.D., Hofmann, N.K., Vance, J.M., Cuccaro, M.L. et al. (2019) RNA editing alterations in a multi-ethnic Alzheimer disease cohort converge on immune and endocytic molecular pathways. Hum. Mol. Genet., 28, 3053–3061.

3. Eisenberg, E. and Levanon, E.Y. (2018) A-to-I RNA editing—immune protector and transcriptome diversifier. Nat. Rev. Genet., 19, 473–490.

4. Lomeli, H., Mosbacher, J., Melcher, T., Hoger, T., Kuner, T., Monyer, H., Higuchi, M., Bach, A. and Seeburg, P.H. (1994) Control of kinetic properties of AMPA receptor channels by nuclear RNA editing. Science, 266, 1709–1713.

5. Wu, S., Yang, M., Kim, P. and Zhou, X. (2021) ADeditome provides the genomic landscape of A-to-I RNA editing in Alzheimer’s disease. Brief. Bioinform.

6. Khermesh, K., D’Erchia, A.M., Barak, M., Annese, A., Wachtel, C., Levanon, E.Y., Picardi, E. and Eisenberg, E. (2016) Reduced levels of protein recoding by A-to-I RNA editing in Alzheimer’s disease. RNA, 22, 290–302.

7. Ma, Y., Dammer, E.B., Felsky, D., Duong, D.M., Klein, H.-U., White, C.C., Zhou, M., Logsdon, B.A., McCabe, C. and Xu, J. (2021) Atlas of RNA editing events affecting protein expression in aged and Alzheimer’s disease human brain tissue. Nat. Commun., 12, 1–16.

8. Pozdyshev, D.V., Zharikova, A.A., Medvedeva, M.V. and Muronetz, V.I. (2021) Differential Analysis of A-to-I mRNA Edited Sites in Parkinson’s Disease. Genes, 13, 14.

9. Gardner, O.K., Van Booven, D., Wang, L., Gu, T., Hofmann, N.K., Whitehead, P.L., Nuytemans, K., Hamilton-Nelson, K.L., Adams, L.D. and Starks, T.D. (2022) Genetic architecture of RNA editing regulation in Alzheimer’s disease across diverse ancestral populations. Hum. Mol. Genet.

10. Park, E., Jiang, Y., Hao, L., Hui, J. and Xing, Y. (2021) Genetic variation and microRNA targeting of A-to-I RNA editing fine tune human tissue transcriptomes. Genome Biol., 22, 1–28.

11. Hodes, R.J. and Buckholtz, N. (2016). Taylor & Francis.

12. Marek, K., Jennings, D., Lasch, S., Siderowf, A., Tanner, C., Simuni, T., Coffey, C., Kieburtz, K., Flagg, E. and Chowdhury, S. (2011) The Parkinson progression marker initiative (PPMI). Prog. Neurobiol., 95, 629–635.

13. Marek, K., Chowdhury, S., Siderowf, A., Lasch, S., Coffey, C.S., Caspell-Garcia, C., Simuni, T., Jennings, D., Tanner, C.M. and Trojanowski, J.Q. (2018) The Parkinson’s progression markers initiative (PPMI)–establishing a PD biomarker cohort. Annals of clinical and translational neurology, 5, 1460–1477.

14. Quinones-Valdez, G., Tran, S.S., Jun, H.-I., Bahn, J.H., Yang, E.-W., Zhan, L., Brümmer, A., Wei, X., Van Nostrand, E.L. and Pratt, G.A. (2019) Regulation of RNA editing by RNA-binding proteins in human cells. Commun. Biol, 2, 1–14.

15. Licht, K., Kapoor, U., Amman, F., Picardi, E., Martin, D., Bajad, P. and Jantsch, M.F. (2019) A high resolution A-to-I editing map in the mouse identifies editing events controlled by pre-mRNA splicing. Genome Res., 29, 1453–1463.

16. Kume, H., Hino, K., Galipon, J. and Ui-Tei, K. (2014) A-to-I editing in the miRNA seed region regulates target mRNA selection and silencing efficiency. Nucleic Acids Res., 42, 10050–10060.

17. Dobin, A., Davis, C.A., Schlesinger, F., Drenkow, J., Zaleski, C., Jha, S., Batut, P., Chaisson, M. and Gingeras, T.R. (2013) STAR: ultrafast universal RNA-seq aligner. Bioinformatics, 29, 15–21.

18. Frankish, A., Diekhans, M., Jungreis, I., Lagarde, J., Loveland, J.E., Mudge, J.M., Sisu, C., Wright, J.C., Armstrong, J. and Barnes, I. (2021) GENCODE 2021. Nucleic Acids Res., 49, D916–D923.

19. Giudice, C.L., Tangaro, M.A., Pesole, G. and Picardi, E. (2020) Investigating RNA editing in deep transcriptome datasets with REDItools and REDIportal. Nat. Protoc., 15, 1098–1131.

20. Mansi, L., Tangaro, M.A., Lo Giudice, C., Flati, T., Kopel, E., Schaffer, A.A., Castrignanò, T., Chillemi, G., Pesole, G. and Picardi, E. (2021) REDIportal: millions of novel A-to-I RNA editing events from thousands of RNAseq experiments. Nucleic Acids Res., 49, D1012–1019.

21. Mailman, M.D., Feolo, M., Jin, Y., Kimura, M., Tryka, K., Bagoutdinov, R., Hao, L., Kiang, A., Paschall, J. and Phan, L. (2007) The NCBI dbGaP database of genotypes and phenotypes. Nat. Genet., 39, 1181–1186.

22. Tryka, K.A., Hao, L., Sturcke, A., Jin, Y., Wang, Z.Y., Ziyabari, L., Lee, M., Popova, N., Sharopova, N. and Kimura, M. (2014) NCBI’s Database of Genotypes and Phenotypes: dbGaP. Nucleic Acids Res., 42, D975–D979.

23. Hinrichs, A.S., Karolchik, D., Baertsch, R., Barber, G.P., Bejerano, G., Clawson, H., Diekhans, M., Furey, T.S., Harte, R.A. and Hsu, F. (2006) The UCSC Genome Browser Database: update 2006. Nucleic Acids Res., D590–D598.

24. Consortium, G. (2015) The Genotype-Tissue Expression (GTEx) pilot analysis: multitissue gene regulation in humans. Science, 348, 648–660.

25. Li, B. and Dewey, C.N. (2011) RSEM: accurate transcript quantification from RNA-Seq data with or without a reference genome. BMC Bioinformatics, 12, 1–16.

26. Kahles, A., Ong, C.S., Zhong, Y. and Rätsch, G. (2016) SplAdder: identification, quantification and testing of alternative splicing events from RNA-Seq data. Bioinformatics, 32, 1840–1847.

27. Li, H. (2013) Aligning sequence reads, clone sequences and assembly contigs with BWA-MEM. arXiv preprint arXiv:1303.3997.

28. Poplin, R., Ruano-Rubio, V., DePristo, M.A., Fennell, T.J., Carneiro, M.O., Van der Auwera, G.A., Kling, D.E., Gauthier, L.D., Levy-Moonshine, A. and Roazen, D. (2018) Scaling accurate genetic variant discovery to tens of thousands of samples. BioRxiv, 201178.

29. Consortium, T.G. (2015) The Genotype-Tissue Expression (GTEx) pilot analysis: Multitissue gene regulation in humans. Science, 348.

30. Yang, Y., Zhang, Q., Miao, Y.-R., Yang, J., Yang, W., Yu, F., Wang, D., Guo, A.-Y. and Gong, J. (2020) SNP2APA: a database for evaluating effects of genetic variants on alternative polyadenylation in human cancers. Nucleic Acids Res., 48, D226–D232.

31. Price, A.L., Patterson, N.J., Plenge, R.M., Weinblatt, M.E., Shadick, N.A. and Reich, D. (2006) Principal components analysis corrects for stratification in genome-wide association studies. Nat. Genet.

32. Stegle, O., Parts, L., Durbin, R., Winn, J. and Regev, A. (2010) A Bayesian Framework to Account for Complex Non-Genetic Factors in Gene Expression Levels Greatly Increases Power in eQTL Studies. PLoS Comput. Biol., 6.

33. Shabalin, A.A. (2012) Matrix eQTL: ultra fast eQTL analysis via large matrix operations. Bioinformatics, 28, 1353–1358.

34. Li, J.-H., Liu, S., Zhou, H., Qu, L.-H. and Yang, J.-H. (2014) starBase v2. 0: decoding miRNA-ceRNA, miRNA-ncRNA and protein–RNA interaction networks from large-scale CLIP-Seq data. Nucleic Acids Res., 42, D92–D97.

35. Bahn, J.H., Ahn, J., Lin, X., Zhang, Q., Lee, J.-H., Civelek, M. and Xiao, X. (2015) Genomic analysis of ADAR1 binding and its involvement in multiple RNA processing pathways. Nat. Commun., 6, 1–13.

36. Schmidt, E.M., Zhang, J., Zhou, W., Chen, J., Mohlke, K.L., Chen, Y.E. and Willer, C.J. (2015) GREGOR: evaluating global enrichment of trait-associated variants in epigenomic features using a systematic, data-driven approach. Bioinformatics, 31, 2601–2606.

37. Teng, X., Chen, X., Xue, H., Tang, Y., Zhang, P., Kang, Q., Hao, Y., Chen, R., Zhao, Y. and He, S. (2020) NPInter v4. 0: an integrated database of ncRNA interactions. Nucleic Acids Res., 48, D160–D165.

38. Lin, Y., Liu, T., Cui, T., Wang, Z., Zhang, Y., Tan, P, Huang, Y., Yu, J. and Wang, D. (2020) RNAInter in 2020: RNA interactome repository with increased coverage and annotation. Nucleic Acids Res., 48, D189–D197.

39. Freund, E.C., Sapiro, A.L., Li, Q., Linder, S., Moresco, J.J., Yates III, J.R. and Li, J.B. (2020) Unbiased Identification of trans Regulators of ADAR and A-to-I RNA Editing. Cell Rep., 31, 107656.

40. Lambert, S.A., Jolma, A., Campitelli, L.F., Das, P.K., Yin, Y., Albu, M., Chen, X., Taipale, J., Hughes, T.R. and Weirauch, M.T. (2018) The human transcription factors. Cell, 172, 650–665.

41. Coetzee, S.G., Coetzee, G.A. and Hazelett, D.J. (2015) motifbreakR: an R/Bioconductor package for predicting variant effects at transcription factor binding sites. Bioinformatics, 31, 3847–3849.

42. Liu, X., Li, C., Mou, C., Dong, Y. and Tu, Y. (2020) dbNSFP v4: a comprehensive database of transcript-specific functional predictions and annotations for human nonsynonymous and splice-site SNVs. Genome Med., 12, 1–8.

43. Lin, S.H., Brown, D.W. and Machiela, M.J. (2020) LDtrait: An Online Tool for Identifying Published Phenotype Associations in Linkage Disequilibrium. Cancer Res., 80, canres.0985.2020.

44. Landrum, M.J., Lee, J.M., Benson, M., Brown, G., Chao, C., Chitipiralla, S., Gu, B., Hart, J., Hoffman, D. and Hoover, J. (2016) ClinVar: public archive of interpretations of clinically relevant variants. Nucleic Acids Res., 44, D862–D868.

45. MacKinnon, D.P., Fairchild, A.J. and Fritz, M.S. (2007) Mediation analysis. Annu. Rev. Psychol., 58, 593.

46. Purcell, S., Neale, B., Todd-Brown, K., Thomas, L., Ferreira, M.A., Bender, D., Maller, J., Sklar, P., De Bakker, P.I. and Daly, M.J. (2007) PLINK: a tool set for whole-genome association and population-based linkage analyses. The American journal of human genetics, 81, 559–575.

47. Kuleshov, M.V., Jones, M.R., Rouillard, A.D., Fernandez, N.F., Duan, Q., Wang, Z., Koplev, S., Jenkins, S.L., Jagodnik, K.M. and Lachmann, A. (2016) Enrichr: a comprehensive gene set enrichment analysis web server 2016 update. Nucleic Acids Res., 44, W90–97.

48. Rangel-Barajas, C., Coronel, I. and Florán, B. (2015) Dopamine receptors and neurodegeneration. Aging and disease, 6, 349.

49. Nobili, A., Latagliata, E.C., Viscomi, M.T., Cavallucci, V., Cutuli, D., Giacovazzo, G., Krashia, P., Rizzo, F.R., Marino, R. and Federici, M. (2017) Dopamine neuronal loss contributes to memory and reward dysfunction in a model of Alzheimer’s disease. Nat. Commun., 8, 1–14.

50. Marambaud, P., Dreses-Werringloer, U. and Vingtdeux, V. (2009) Calcium signaling in neurodegeneration. Mol. Neurodegener., 4, 1–15.

51. Kong, X., Yuan, Z. and Cheng, J. (2017) The function of NOD-like receptors in central nervous system diseases. J. Neurosci. Res., 95, 1565–1573.

52. Ferencz, B., Karlsson, S. and Kalpouzos, G. (2012) Promising genetic biomarkers of preclinical Alzheimer’s disease: the influence of APOE and TOMM40 on brain integrity. International Journal of Alzheimer’s Disease, 2012.

53. Mise, A., Yoshino, Y., Yamazaki, K., Ozaki, Y., Sao, T., Yoshida, T., Mori, T., Mori, Y., Ochi, S. and Iga, J.-i. (2017) TOMM40 and APOE gene expression and cognitive decline in Japanese Alzheimer’s disease subjects. J. Alzheimer’s Dis., 60, 1107–1117.

54. (2018) The NHGRI-EBI GWAS Catalog of published genome-wide association studies, targeted arrays and summary statistics 2019. %J Nucleic Acids Research.

55. Barbarino, J.M., Whirl - Carrillo, M., Altman, R.B. and Klein, T.E. (2018) PharmGKB: a worldwide resource for pharmacogenomic information. Wiley Interdisciplinary Reviews: Systems Biology and Medicine, 10, e1417.

56. Hong, H., An, O., Chan, T.H., Ng, V.H., Kwok, H.S., Lin, J.S., Qi, L., Han, J., Tay, D.J. and Tang, S.J. (2018) Bidirectional regulation of adenosine-to-inosine (A-to-I) RNA editing by DEAH box helicase 9 (DHX9) in cancer. Nucleic Acids Res., 46, 7953–7969.

57. Wang, Z., Zhao, Y., Xu, N., Zhang, S., Wang, S., Mao, Y., Zhu, Y., Li, B., Jiang, Y. and Tan, Y. (2019) NEAT1 regulates neuroglial cell mediating Aβ clearance via the epigenetic regulation of endocytosis-related genes expression. Cell. Mol. Life Sci., 76, 3005–3018.

58. An, H., Williams, N.G. and Shelkovnikova, T.A. (2018) NEAT1 and paraspeckles in neurodegenerative diseases: A missing lnc found? Non-coding RNA research, 3, 243–252.

59. Qiang, J.K., Wong, Y.C., Siderowf, A., Hurtig, H.I., Xie, S.X., Lee, V.M.Y., Trojanowski, J.Q., Yearout, D., B. Leverenz, J. and Montine, T.J. (2013) Plasma apolipoprotein A1 as a biomarker for Parkinson disease. Ann. Neurol., 74, 119–127.

60. Aktaş, T., Avşar Ilik, i., Maticzka, D., Bhardwaj, V., Pessoa Rodrigues, C., Mittler, G., Manke, T., Backofen, R. and Akhtar, A. (2017) DHX9 suppresses RNA processing defects originating from the Alu invasion of the human genome. Nature, 544, 115–119.

61. Stellos, K., Gatsiou, A., Stamatelopoulos, K., Matic, L.P., John, D., Lunella, F.F., Jaé, N., Rossbach, O., Amrhein, C. and Sigala, F. (2016) Adenosine-to-inosine RNA editing controls cathepsin S expression in atherosclerosis by enabling HuR-mediated post-transcriptional regulation. Nat. Med., 22, 1140–1150.

62. Jiang, F., Liu, X., Wang, X., Hu, J., Chang, S. and Cui, X. (2022) LncRNA FGD5-AS1 accelerates intracerebral hemorrhage injury in mice by adsorbing miR-6838-5p to target VEGFA. Brain Res., 1776, 147751.

63. Xu, F., Na, L., Li, Y. and Chen, L. (2020) RETRACTED ARTICLE: Roles of the PI3K/AKT/mTOR signalling pathways in neurodegenerative diseases and tumours. Cell & bioscience, 10, 1–12.

64. Nishikura, K. (2016) A-to-I editing of coding and non-coding RNAs by ADARs. Nat. Rev. Mol. Cell Biol., 17, 83–96.

65. Tran, S.S., Jun, H.-I., Bahn, J.H., Azghadi, A., Ramaswami, G., Van Nostrand, E.L., Nguyen, T.B., Hsiao, Y.-H.E., Lee, C. and Pratt, G.A. (2019) Widespread RNA editing dysregulation in brains from autistic individuals. Nat. Neurosci., 22, 25–36.

66. Storey, J.D. and Tibshirani, R. (2003) Statistical significance for genomewide studies. Proceedings of the National Academy of Sciences, 100, 9440–9445.

67. Gellersen, H.M., Guo, C.C., O’Callaghan, C., Tan, R.H., Sami, S. and Hornberger, M. (2017) Cerebellar atrophy in neurodegeneration—a meta-analysis. J. Neurol. Neurosurg. Psychiatry, 88, 780–788.

68. Azam, S., Haque, M.E., Jakaria, M., Jo, S.-H., Kim, I.-S. and Choi, D.-K. (2020) G-protein-coupled receptors in CNS: a potential therapeutic target for intervention in neurodegenerative disorders and associated cognitive deficits. Cells, 9, 506.

69. Hashemi-Firouzi, N., Komaki, A., Asl, S.S. and Shahidi, S. (2017) The effects of the 5-HT7 receptor on hippocampal long-term potentiation and apoptosis in a rat model of Alzheimer’s disease. Brain Res. Bull., 135, 85–91.

70. Matamoros, A.J. and Baas, P.W. (2016) Microtubules in health and degenerative disease of the nervous system. Brain Res. Bull., 126, 217–225.

71. Ng, B., White, C.C., Klein, H.-U., Sieberts, S.K., McCabe, C., Patrick, E., Xu, J., Yu, L., Gaiteri, C. and Bennett, D.A. (2017) An xQTL map integrates the genetic architecture of the human brain’s transcriptome and epigenome. Nat. Neurosci., 20, 1418–1426.

72. Aykaç, A. and Sehirli, A.O. (2020) The role of the SLC transporters protein in the neurodegenerative disorders. Clinical Psychopharmacology and Neuroscience, 18, 174.

73. Wolozin, B. and Ivanov, P. (2019) Stress granules and neurodegeneration. Nat. Rev. Neurosci., 20, 649–666.

74. Kong, W., Mou, X., Liu, Q., Chen, Z., Vanderburg, C.R., Rogers, J.T. and Huang, X. (2009) Independent component analysis of Alzheimer’s DNA microarray gene expression data. Mol. Neurodegener., 4, 1–14.

75. Cui, C., Sun, J., Pawitan, Y., Piehl, F., Chen, H., Ingre, C., Wirdefeldt, K., Evans, M., Andersson, J. and Carrero, J.-J. (2020) Creatinine and C-reactive protein in amyotrophic lateral sclerosis, multiple sclerosis and Parkinson’s disease. Brain communications, 2, fcaa152.

76. Yang, J., Bennett, B.D., Luo, S., Inoue, K., Grimm, S.A., Schroth, G.P., Bushel, P.R., Kinyamu, H.K. and Archer, T.K. (2015) LIN28A modulates splicing and gene expression programs in breast cancer cells. Mol. Cell. Biol., 35, 3225–3243.

77. Paz, I., Kosti, I., Ares Jr, M., Cline, M. and Mandel-Gutfreund, Y. (2014) RBPmap: a web server for mapping binding sites of RNA-binding proteins. Nucleic Acids Res., 42, W361–W367.

78. Voronin, M.V., Kadnikov, I.A., Zainullina, L.F., Logvinov, I.O., Verbovaya, E.R., Antipova, T.A., Vakhitova, Y.V. and Seredenin, S.B. (2021) Neuroprotective Properties of Quinone Reductase 2 Inhibitor M-11, a 2-Mercaptobenzimidazole Derivative. Int. J. Mol. Sci., 22, 13061.

79. Wickramasinghe, V.O., Gonzàlez-Porta, M., Perera, D., Bartolozzi, A.R., Sibley, C.R., Hallegger, M., Ule, J., Marioni, J.C. and Venkitaraman, A.R. (2015) Regulation of constitutive and alternative mRNA splicing across the human transcriptome by PRPF8 is determined by 5’splice site strength. Genome Biol., 16, 1–21.

80. Glasser, E., Maji, D., Biancon, G., Puthenpeedikakkal, A.M.K., Cavender, C.E., Tebaldi, T., Jenkins, J.L., Mathews, D.H., Halene, S. and Kielkopf, C.L. (2022) Pre-mRNA splicing factor U2AF2 recognizes distinct conformations of nucleotide variants at the center of the pre-mRNA splice site signal. Nucleic Acids Res., 50, 5299–5312.

81. Yang, E.-W., Bahn, J.H., Hsiao, E.Y.-H., Tan, B.X., Sun, Y., Fu, T., Zhou, B., Van Nostrand, E.L., Pratt, G.A. and Freese, P. (2019) Allele-specific binding of RNA-binding proteins reveals functional genetic variants in the RNA. Nat. Commun., 10, 1–15.

