## Supplementary figures and images for "Genetic control of RNA editing in Neurodegenerative disease"

### Supplementary Figure 1

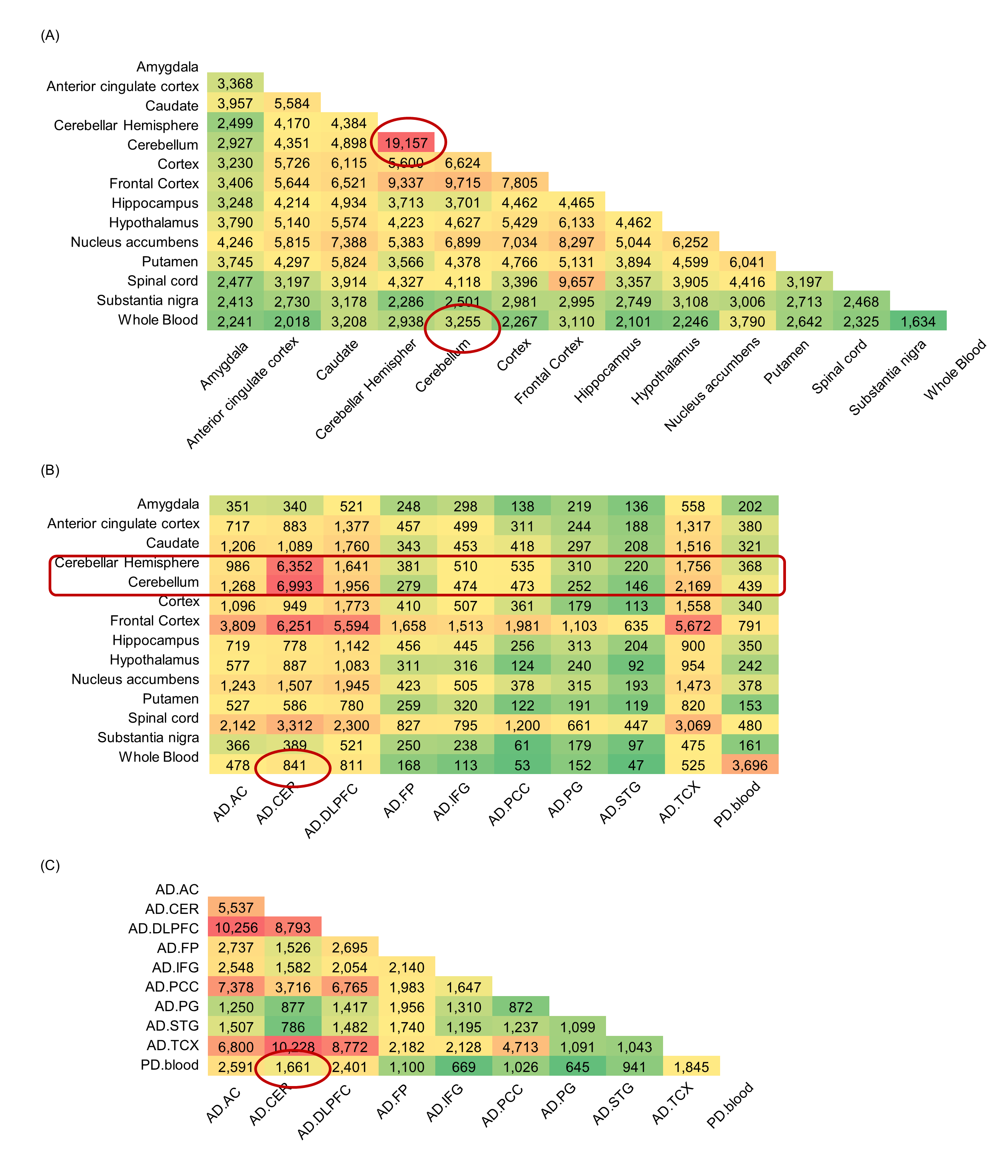

### Supplementary Figure 2

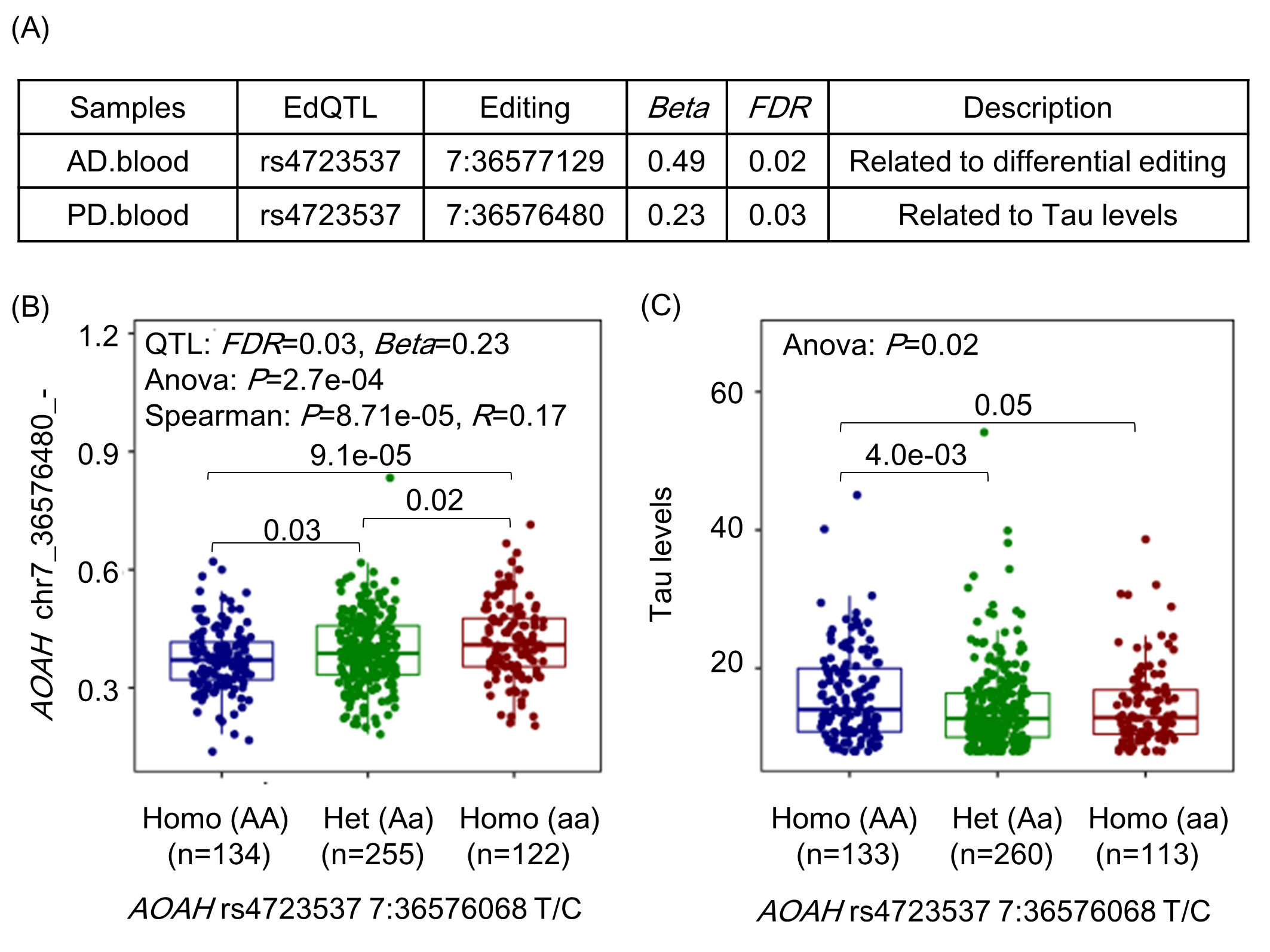
